# AdmirePred: A method for predicting abundant miRNAs in Exosomes

**DOI:** 10.1101/2025.03.19.644072

**Authors:** Akanksha Arora, Gajendra Pal Singh Raghava

**Affiliations:** Department of Computational Biology, Indraprastha Institute of Information Technology, Okhla Phase 3, New Delhi-110020, India

**Keywords:** Exosome, Extracellular Vesicle, miRNA, Machine learning, Alignment-based, Non-invasive diagnostics, Liquid Biopsy

## Abstract

Non-invasive disease diagnosis is a key application of blood exosomes in liquid biopsy, as they carry diverse biological molecules, including microRNAs (miRNAs) derived from their parent cells. Developing miRNA-based disease biomarkers requires prediction of highly abundant miRNAs in exosomes under normal conditions for establishing a baseline for understanding their physiological roles and disease-specific variations. In this study, we present models for predicting highly abundant miRNAs in exosomes from their nucleotide sequences. The models were trained, tested, and evaluated on a dataset comprising 348 abundant and 349 non-abundant miRNAs. Initially, we applied alignment-based approaches, such as motif and similarity searches, but these methods yielded poor coverage. We then explored alignment-free approaches, particularly machine learning models leveraging a broad range of features. Our Extra Trees classifier, developed using binary profiles and TF-IDF features, achieved the highest performance with an AUC of 0.77. To further enhance predictive accuracy, we developed a hybrid method that combines machine learning models with alignment-based approaches, achieving an AUC of 0.854 on an independent dataset. To support research in non-invasive diagnostics and therapeutics, we have developed a web server, standalone tool, and Python package for AdmirePred, available at https://webs.iiitd.edu.in/raghava/admirepred/.

**Key points:**

1. miRNA abundant in blood exosomes are promising biomarkers for liquid biopsy
2. Classification of abundant and non-abundant miRNA in healthy individuals
3. A hybrid method that combine alignment based and alignment free approach
4. Prediction of miRNAs that are highly expressed in blood exosomes
5. A web server, a python package, and a standalone tool have been created

**Author’s Biography:**

1. Akanksha Arora is currently pursuing a Ph.D. in Computational Biology at Department of Computational Biology, Indraprastha Institute of Information Technology, New Delhi, India.
2. Gajendra P. S. Raghava is currently working as a Professor and Head of Department of Computational Biology, Indraprastha Institute of Information Technology, New Delhi, India.

## 1. Introduction

Exosomes are small extracellular vesicles, typically 30-150 nm in size, are secreted by nearly all cell types and carry a cargo of proteins, lipids, and nucleic acids that reflect the physiological and pathological state of their parent cells. Their stability in bodily fluids such as blood, urine, and saliva makes them ideal candidates for minimally invasive diagnostic and prognostic applications [1–4]. Exosomes are gaining significant attention in the field of liquid biopsy as promising non-invasive biomarkers for various diseases, including cancer, cardiovascular conditions, and neurodegenerative disorders [5–8]. Recent advancements in isolation and analytical techniques have enhanced the ability to harness exosomes for early disease detection, monitoring treatment response, and understanding disease mechanisms. As research progresses, exosomes are increasingly recognized for their potential to revolutionize personalized medicine by providing real-time insights into disease progression and therapeutic efficacy [9,10].

These vesicles carry a diverse array of molecular cargo, including mRNA, miRNA, lipids, proteins, and DNA. Among these, microRNAs (miRNAs) have garnered significant attention due to their regulatory influence on gene expression and their potential as diagnostic and prognostic biomarkers for various diseases [11–14]. These small non-coding RNAs (typically 20–22 nucleotides in length) modulate post-transcriptional gene expression by binding to target mRNAs, leading to their degradation or translational repression. MiRNAs abundant in exosomes are particularly significant as they regulate crucial biological pathways involved in cell proliferation, differentiation, apoptosis, and immune response modulation. Their stability in bodily fluids and ability to reflect disease-specific alterations make them valuable biomarkers for liquid biopsies, offering diagnostic, prognostic, and therapeutic insights into conditions such as cancer, cardiovascular diseases, and neurodegenerative disorders.

While exosomes have emerged as a focal point in liquid biopsy research, significant efforts have also been directed toward understanding the molecules that are localized in exosomes as their presence in exosomes or other cellular compartments can influence their biological roles and utility as biomarkers [15–18]. Earlier we developed a method – EmiRPred, that classifies exosomal and non-exosomal miRNA based on the dataset collected from RNALocate, where subcellular locations for miRNA are given [18,19]. However, earlier we did not take miRNA expression into account and classified them as exosomal and non-exosomal. miRNAs exist in multiple subcellular locations, including the cytoplasm, nucleus, mitochondria, extracellular space (both vesicular and non-vesicular forms), and ribonucleoprotein complexes, but their expression levels vary amongst the locations [20,21]. Instead of using the binary approach of whether an miRNA is observed in exosome or not, we used expression-based approach that identifies miRNA that are abundant in exosomes. In this study, we have made a systematic effort to identify and predict the miRNA that are abundantly present in exosomes of normal subjects by taking their expression values from various studies. We believe that discovering which miRNA are present inside exosomes under normal conditions will help the scientists discover new biomarkers for a number of diseases, as well as enabling more precise and non-invasive disease monitoring [22,23].

Here, we present AdmirePred, a computational method designed to predict abundant miRNAs in exosomes. In this method, we have integrated two types of techniques including alignment-based approaches and AI-based approaches developed on a set of features. The method is trained and validated on experimentally verified miRNA datasets extracted from EVmiRNA and GEO datasets [24,25]. The architecture of this study is shown in Figure 1.

**Figure 1:**
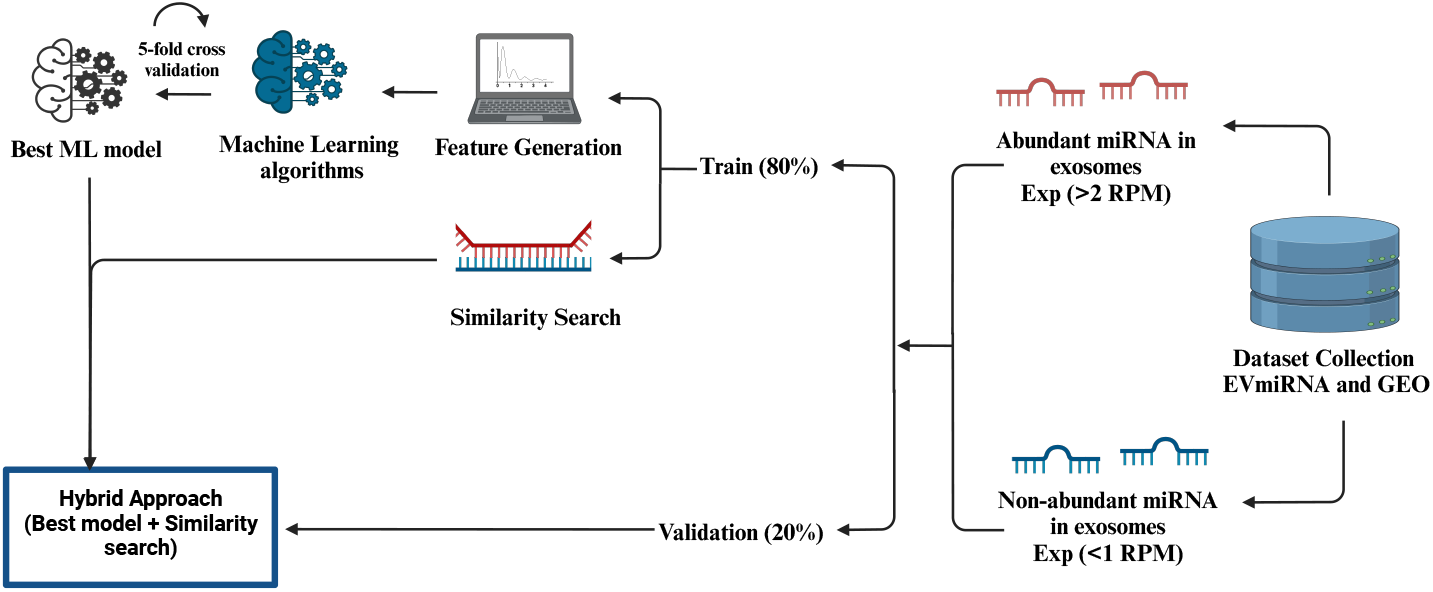
Architecture of AdmirePred

## 2. Methods

### 2.1. Data Collection and Preprocessing

We collected the data from EVmiRNA for blood exosomes (serum and plasma) for normal human subjects (n=60), and average expressions were taken into account. The miRNAs having average expression in Reads Per Kilobase Million (RPKM) > 2 were assigned as abundant miRNA in exosomes whereas miRNA having average expression in RPKM < 1 were assigned as non-abundant miRNA. In this study, we refer abundant miRNAs in exosomes as exosomal miRNAs and non-abundant miRNAs as non-exosomal miRNAs. This data was then validated with another GEO study GSE270497 where the small RNA sequencing of plasma exosomes was done for both breast cancer and normal human subjects. We took the expression data for normal subjects into account to verify our data. Altogether, after obtaining the data for miRNA sequences, and removing the duplicates, we were able to get about 348 abundant and 349 non-abundant sequences, where the sequences ranged from 16 to 25 nucleotides.

### 2.2. Motif-Search

We utilized the MERCI (Motif Emerging with Classes Identification) tool to identify motifs within miRNA sequences that are abundantly present in exosomes [26]. The motifs were discovered using the training dataset and then those motifs were searched within an independent validation set. The MERCI software allows for the selection of motifs that are exclusively present in the positive class (miRNA abundantly present in exosomes) within the training data. Furthermore, the software provides several customization options, including the ability to analyze motif frequency and detect motifs with or without gaps [16].

### 2.3. Similarity Search

In this study, we employed the blastn-short tool to annotate miRNA sequences by comparing their similarity to known abundant and non-abundant miRNA sequences in exosomes [27]. Initially, a database was created using the “makeblastdb” command, incorporating miRNA sequences from the training dataset. To generate results for the training set, self-hits were excluded, and only the top hit, after removing self-hits, was considered. For the independent validation set, the first hit was used to evaluate results across different e-value thresholds. This approach has been previously utilized in earlier studies [28–30].

### 2.4. ML-based classification methods

#### 2.4.1. Feature Generation

To build a prediction model for distinguishing between abundant and non-abundant miRNA in exosomes, we employed various techniques to extract relevant features from the miRNA sequences. These techniques are outlined below:

a) ***Nucleotide Composition:*** We employed Nfeature to calculate a diverse set of composition-based features from miRNA sequences, such as nucleotide composition of a sequence as well as its reverse complementary sequence for k-mers = 1,2,3, and 4 (nucleotide sequences of length k) [31].
b) ***Nucleotide-weighted frequency:*** We extracted weighted frequency of nucleotides using Term Frequency-Inverse Document Frequency (TF-IDF), a statistical technique that determines the significance of a term within a document in relation to a larger collection of documents. In our context, “terms” correspond to **k-mers**, and a “document” represents an miRNA sequence [32]. The Term Frequency (TF) measures how often a specific k-mer appears within a sequence, normalized by the total number of k-mers in that sequence. The Inverse Document Frequency (IDF) quantifies the uniqueness of a k-mer by calculating the logarithmic inverse of the fraction of sequences in the dataset that contain the k-mer. By combining these values, TF-IDF assigns higher importance to k-mers that frequently appear in specific sequences but are relatively rare across the dataset, making them more informative and representative of the sequence’s characteristics. These metrics are further explained in Equations 1-3.

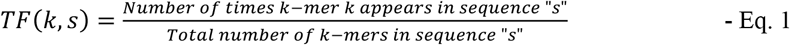

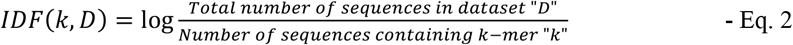

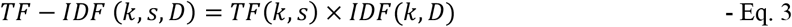

In these equations, *k* represents the k-mer, *s* denotes a sequence, and *D* stands for the entire dataset of sequences. TF-IDF features were computed for both the original and reverse complementary sequences, covering k-mer ranges from (1,1) to (1,7).
c) ***Distance Distribution of Nucleotides:*** We calculated distance distribution of different nucleotide types in miRNA sequences using Nfeature tool. Distance-based features help uncover hidden sequence patterns that might not be evident from simple nucleotide composition (like k-mer frequencies) consider direct compositions but often ignore spatial relationships between nucleotides. However, in biological sequences like miRNAs, the relative positioning of nucleotides might play a crucial role in their structural stability, functional interactions, and regulatory activity.
d) ***Nucleotide Repeat Index:*** The Nucleotide Repeats Index (NRI) concisely captures the extent of consecutive runs of each nucleotide (A, C, G, U) in a sequence, turning those repeats into a simple numeric descriptor. By highlighting how nucleotides cluster or disperse, NRI aids in detecting patterns which can influence RNA folding and gene regulation.
e) ***Entropy:*** The Shannon Entropy feature gauges the randomness or unpredictability of nucleotides in a given sequence. It reflects how uniformly the four nucleotides (A, C, G, U) are distributed—sequences dominated by one or two nucleotides have lower entropy, while more balanced sequences show higher entropy. This measure helps distinguish sequences based on complexity and can highlight potential regulatory or structurally significant regions where nucleotide diversity (or lack thereof) matters. We calculated both sequence level and nucleotide level Shannon Entropy for our dataset. The sequence level Shannon Entropy produces one overall entropy value for the entire sequence, whereas the nucleotide level Shannon entropy breaks the calculation down to yield a 4-value vector, capturing how each nucleotide’s distribution contributes to the entropy.
f) ***Correlation:*** We computed correlation-based descriptors to capture positional relationships and physicochemical similarities within miRNA sequences. We computed a) Autocorrelation which measures how the same property (e.g., a certain physicochemical index) at different positions correlates over a given “lag” (distance); b) Cross-correlation which evaluates how two different properties correlate at positions separated by a specific lag; c) Auto-cross correlation: A hybrid approach that merges the ideas of both autocorrelation (same property) and cross-correlation (different properties) in a single descriptor.
g) ***Pseudo Composition:*** Compared with the conventional nucleotide composition, pseudo nucleotide compositions have the advantage of converting RNA sequences of various lengths to a fixed-length digital vector to enable sequence comparison, while at the same time keeping the long-range sequence order information. Along with pseudo composition, we also computed two subtypes namely serial correlation pseudo composition which captures the dependency between consecutive dinucleotides in a sequence, and parallel correlation pseudo composition which computes multiple correlations simultaneously across different sequence positions.
h) ***Binary Profiles:*** We generated binary profiles, also known as one-hot encoding features, to represent nucleotide sequences as binary vectors. To accommodate sequences of varying lengths, we applied a padding technique to standardize their length. As the longest miRNA sequence in our study is 25 nucleotides, all sequences were extended to this length by appending ‘X’ to the sequences until they have a length of 25 nucleotides. For instance, a sequence such as “AGTAGCATCGUUAAGTUCCAAU” which has a length of 22, was padded to “AGTAGCATCGUUAAGTUCCAAUXXX” to reach the required length of 25. Once padding was applied, the sequences were transformed into binary profiles using one-hot encoding.

#### 2.4.2. Prediction Models

We employed multiple machine learning (ML) algorithms from Scikit-learn to classify abundant and non-abundant miRNA sequences in exosomes. Decision Tree (DT) splits data based on feature thresholds to form decision rules, Random Forest (RF) builds multiple decision trees and averages their predictions for better accuracy, Logistic Regression (LR) estimates probabilities using a logistic function, K-Nearest Neighbors (KNN) classifies based on the majority class of k nearest data points, Gaussian Naïve Bayes (GNB) applies Bayes’ theorem with a Gaussian distribution, Support Vector Classifier (SVC) finds the optimal hyperplane to separate classes, Extreme Gradient Boosting (XGB) improves predictions through iterative gradient-boosted decision trees, and Extra Trees Classifier (ET) enhances RF by adding randomness to feature selection and splits, improving robustness [33–38]. Prediction models were developed using these algorithms based on various feature sets, as described in Section 2.3.1.

#### 2.4.3. Cross-validation and performance metrics

The dataset, comprising 697 sequences, was split in an 80:20 ratio, with 80% allocated for training and 20% for validation. To assess the performance of the ML models, a five-fold cross-validation approach was applied to the training set, while the remaining 20% was kept separate and unseen during training the models. In five-fold cross-validation, the training data is divided into five sets, with four sets used for training and one for internal validation, rotating this process five times so that each set is used as a validation set once. The models were evaluated using both threshold-dependent and threshold-independent performance metrics. The evaluation criteria included the calculation of specificity, sensitivity, accuracy Matthews correlation coefficient (MCC), and the Area Under the Receiver Operating Characteristic Curve (AUROC). AUROC is independent of thresholds, whereas the other metrics rely on threshold values. Sensitivity, specificity, and MCC were optimized to identify the best threshold. These evaluation metrics have been widely utilized in previous research to assess model performance and are shown in Equations 4-7 [39–43].

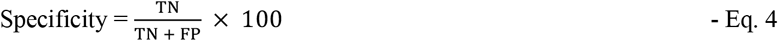

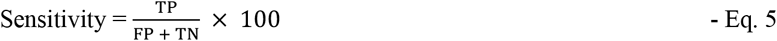

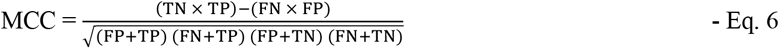

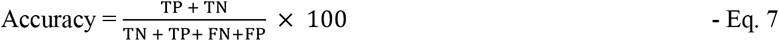

Where FP, TP, FN, and TN are false positive, true positive, false negative, and true negative, respectively.

### 2.5. Ensemble method

To enhance the predictive performance of our top-performing model, developed using the optimal feature set, we developed a method that integrates different approaches used in this study. This method employs a weighted scoring system that combines three techniques: (i) a motif-based analysis, (ii) similarity search via BLAST, and (iii) ML predictions. In this scoring scheme, an miRNA sequence is awarded +0.5 points if an exosomal motif is detected and 0 points if no such motif is found and added to the predicted probabilities generated by the best-performing ML model using the predict_proba() function from scikit-learn, which provides the likelihood of a sequence belonging to a particular class rather than a simple binary classification [44]. Similarly +0.5 points are assigned if the sequence is identified as similar to an exosomal (abundant in exosomes) miRNA through BLAST analysis and added to the probabilities generated by best performing ML model. The combined score of ML and motif-search varies from 0 to 1.5, and the combined score for ML and blast-search varies from 0 to 1.5 as well. These are defined in equations 8 and 9. Based on these cumulative scores, sequences were classified as either exosomal or non-exosomal. This hybrid or ensemble approach has been previously applied in various studies [16,28].

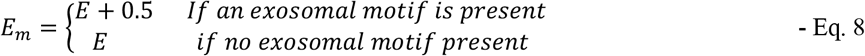

Here, E = Prediction probability score acquired from best performing ML model and *E*_*m*_ = Score obtained after the scores from the motif-based approach are added

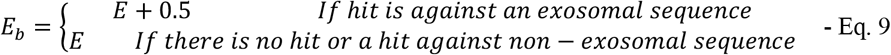

Here, *E*_*b*_= Final score obtained by adding scores from the best performing ML model and BLAST-based approach ranging from 0 to 1.5

## 3. Results

In this study, we used various techniques to predict abundant miRNA in exosomes, which can be broadly classified into three categories: (i) Alignment-based methods, (ii) AI-based methods, and (iii) ensemble methods. The alignment-based methods include motif identification using the MERCI tool and similarity search through BLAST. The AI-based methods involve the application of ML models on a diverse range of features and their combinations. To maximize the advantages of both alignment-based and AI-based approaches, we developed an ensemble strategy that effectively integrates their strengths.

### 3.1 Motif-Search

In motif-search using MERCI, it was observed that there was low coverage and a high error rate with a coverage of only 8 sequences in the independent validation set with 3 wrong predictions for gap = 0. Similarly for gap=1, about 15 sequences were covered with 6 wrong predictions.

### 3.2. Similarity-search using BLAST

In this study, we utilized the blastn-short tool to implement similarity searches against a training dataset comprising abundant and non-abundant miRNA sequences in exosomes, exploring e-values from 10^−6^ to 10^6^ [27]. The optimal performance was observed at e-value 10^−2^, with 24 correct hits and 0 incorrect hits for exosomal (abundant in exosomes) miRNA sequences out of the total of 69 exosomal sequences in the validation set. The e-values below 10^−2^ resulted in insufficient sequence coverage, while values exceeding 10^−2^ led to an increased error rate. The complete results for BLAST for the e-values 10^−6^ to 10^6^ are described in Table 1.

**Table 1:** Number of correct, incorrect, and total hits for training and independent validation sets comprising exosomal (abundant in exosomes) miRNA sequences for e-values ranging from 10^−6^ to 10^6^

|  | Training set |  |  |  | Validation set |  |  |  |
| --- | --- | --- | --- | --- | --- | --- | --- | --- |
| e-value | Correct hits | Incorrect hits | No hits | Total | Correct hits | Incorrect hits | No hits | Total |
| $10^6$ | 196 | 83 | 0 | 279 | 32 | 37 | 0 | 69 |
| $10^5$ | 196 | 83 | 0 | 279 | 32 | 37 | 0 | 69 |
| $10^4$ | 196 | 83 | 0 | 279 | 32 | 37 | 0 | 69 |
| $10^3$ | 196 | 83 | 0 | 279 | 32 | 37 | 0 | 69 |
| $10^2$ | 196 | 83 | 0 | 279 | 32 | 37 | 0 | 69 |
| $10^1$ | 196 | 83 | 0 | 279 | 32 | 37 | 0 | 69 |
| $10^0(1)$ | 167 | 57 | 55 | 279 | 43 | 11 | 15 | 69 |
| $10^{-1}$ | 118 | 31 | 130 | 279 | 34 | 6 | 29 | 69 |
| $10^{-2}$ | 85 | 6 | 188 | 279 | 24 | 0 | 45 | 69 |
| $10^{-3}$ | 61 | 1 | 217 | 279 | 14 | 0 | 55 | 69 |
| $10^{-4}$ | 51 | 1 | 227 | 279 | 13 | 0 | 56 | 69 |
| $10^{-5}$ | 35 | 1 | 243 | 279 | 10 | 0 | 59 | 69 |
| $10^{-6}$ | 14 | 1 | 264 | 279 | 2 | 0 | 67 | 69 |

### 3.3. ML-based classification methods

First, we extracted a comprehensive set of sequence features from miRNA sequences, as described in the Materials & Methods section. Subsequently, we employed various machine learning algorithms to build prediction models, including (LR), Gaussian Naïve Bayes (GNB), K-Nearest Neighbors (KNN), Decision Tree (DT), Support Vector Classifier (SVC), Extreme Gradient Boosting (XGB), Random Forest (RF), and Extra Trees Classifier (ET) models. The results are shown in Supplementary Table S1. The performances of ML-based models developed using various types of features are given below:

a) ***Nucleotide Composition:*** We calculated nucleotide composition of miRNA sequences as well as their reverse complementary sequences for k-mers 1 to 4. The best performing feature among this category was the composition of reverse complementary miRNA sequence for k-mer =3, where the highest AUC of 0.738 on independent validation set was achieved on an SVC model.
b) ***Nucleotide-weighted frequency:*** The best performing composition-based features were TF-IDF for reverse complementary sequences for k-mers 1 and 2 leading to an AUC of 0.744 in the independent validation set for ET model.
c) ***Distance Distribution of Nucleotides:*** We computed distance distribution of nucleotides using Nfeature, and got the highest performance of 0.609 AUC for training set and 0.621 on independent validation set.
d) ***Nucleotide Repeat Index:*** The nucleotide repeat index was calculated as described in the Methods section. The highest AUC observed for these set of features was 0.593 and 0.594 for training and independent validation sets respectively.
e) ***Entropy:*** We computed Shannon Entropy on two levels including sequence level and nucleotide level entropy. The highest AUC achieved for sequence level was 0.561 on independent validation set, whereas for nucleotide level it was 0.594 for independent validation set.
f) ***Correlation:*** We calculated three types of correlations in this study including autocorrelation, auto-cross correlation, and cross correlation. For this, the best performing features were autocorrelation of dinucleotides with AUC of 0.62 for independent validation set.
g) ***Pseudo Composition:*** We computed pseudo composition, serial correlation pseudo composition, parallel correlation pseudo compositions for dipeptides. The highest performing feature was serial correlation pseudo composition with an AUC of 0.708 for training and 0.707 for independent validation set.
h) ***Binary Profiles:*** We calculated binary profiles or one hot encoding features for the miRNA sequences to represent these sequences as binary vectors. These features were one of the best performing features with the highest AUC of 0.75 in the independent validation set.
i) ***Best features:*** We developed the machine learning models by combining the best features in composition-based features and binary profile-based features. The best performing model was Extra Tree classifier resulting in an AUC of 0.763 in training and 0.769 in independent validation set. The results for best features are given in Table 2 and the results for combined best features are shown in Table 3. We also performed a Mann-Whitney test on these features and it was found that 31 features were significantly different (p-value<0.05) from a total of 145 features. The results for Mann-Whitney Test are given in Supplementary Table S2.
j) ***Feature Importance*** We selected the best-performing 20 features using the feature importance in the ET model for alternate dataset as well. About 15 of the 20 were found to be significantly different (p-value<0.05) between the exosomal and non-exosomal classes. In binary profile features, G at 15^th^, G at 19^th^, A at 16^th^, G at 10^th^ and positions were found to be significantly increased in exosomal sequences, whereas, U at 15^th^ position, A at 20^th^, and A at 11^th^ were found to be significantly decreased in exosomal sequences as compared to non-exosomal sequences. In TFIDF features from the range of (1,2) in reverse complement miRNA sequences, it is seen that the occurrence of C, AC, CG, and GA are significantly increased in reverse complementary exosomal sequences, and UU, U, AU, and UA are significantly decreased in reverse complementary exosomal sequences as compared to non-exosomal sequences. The features with their importance are given in Supplementary Table S3, and the comparison between the two classes for top 20 important features are shown in Figure 2.

**Table 2:** The performance of machine learning-based methods developed using binary profile

| Binary profile-based features |  |  |  |  |  |  |  |  |  |  |  |
| --- | --- | --- | --- | --- | --- | --- | --- | --- | --- | --- | --- |
|  |  | Training dataset (cross-validation) |  |  |  |  | Validation/Independent dataset |  |  |  |  |
| Model | Thr | Sens | Spec | Acc | AUC | MCC | Sens | Spec | Acc | AUC | MCC |
| DT | 0.49 | 55.91 | 56.63 | 56.27 | 0.581 | 0.125 | 52.17 | 57.14 | 54.68 | 0.563 | 0.093 |
| RF | 0.51 | 67.38 | 65.59 | 66.49 | 0.747 | 0.330 | 68.12 | 70.00 | 69.06 | 0.754 | 0.381 |
| LR | 0.51 | 63.08 | 63.44 | 63.26 | 0.685 | 0.265 | 66.67 | 54.29 | 60.43 | 0.665 | 0.211 |
| XGB | 0.51 | 62.37 | 61.65 | 62.01 | 0.685 | 0.240 | 68.12 | 52.86 | 60.43 | 0.683 | 0.212 |
| KNN | 0.43 | 67.03 | 63.08 | 65.05 | 0.709 | 0.301 | 76.81 | 57.14 | 66.91 | 0.713 | 0.346 |
| GNB | 0.51 | 62.37 | 63.08 | 62.72 | 0.673 | 0.254 | 71.01 | 57.14 | 64.03 | 0.670 | 0.284 |
| SVC | 0.51 | 64.52 | 64.52 | 64.52 | 0.702 | 0.290 | 79.71 | 57.14 | 68.35 | 0.706 | 0.378 |
| ET | 0.51 | 64.16 | 68.46 | 66.31 | 0.740 | 0.326 | 68.12 | 60.00 | 64.03 | 0.745 | 0.282 |
| Term Frequency- Inverse Document Frequency (TFIDF)<br>on reverse complementary sequence (k-mer range – 1,2) |  |  |  |  |  |  |  |  |  |  |  |
|  |  | Training dataset (cross-validation) |  |  |  |  | Validation/Independent dataset |  |  |  |  |
| Model | Thr | Sens | Spec | Acc | AUC | MCC | Sens | Spec | Acc | AUC | MCC |
| DT | 0.51 | 54.48 | 63.44 | 58.96 | 0.590 | 0.180 | 68.12 | 54.29 | 61.15 | 0.612 | 0.226 |
| RF | 0.51 | 64.88 | 65.59 | 65.23 | 0.701 | 0.305 | 65.22 | 60.00 | 62.59 | 0.695 | 0.252 |
| LR | 0.49 | 62.01 | 59.50 | 60.75 | 0.653 | 0.215 | 68.12 | 54.29 | 61.15 | 0.636 | 0.226 |
| XGB | 0.49 | 62.01 | 62.01 | 62.01 | 0.688 | 0.240 | 66.67 | 52.86 | 59.71 | 0.659 | 0.197 |
| **KNN** | 0.41 | 59.50 | 62.37 | 60.93 | 0.645 | 0.219 | 59.42 | 64.29 | 61.87 | 0.649 | 0.237 |
| **GNB** | 0.49 | 62.72 | 62.37 | 62.55 | 0.658 | 0.251 | 63.77 | 54.29 | 58.99 | 0.645 | 0.181 |
| **SVC** | 0.5 | 65.23 | 65.23 | 65.23 | 0.696 | 0.305 | 62.32 | 57.14 | 59.71 | 0.664 | 0.195 |
| **ET** | 0.5 | 66.31 | 64.16 | 65.23 | 0.718 | 0.305 | 73.49 | 61.43 | 67.63 | 0.744 | 0.356 |

**Table 3:** The performance of machine learning-based models developed using combination of binary profile and reverse complement TFIDF

| Best features combined –<br>Binary-profile based features + Reverse complement TFIDF (1,2) |  |  |  |  |  |  |  |  |  |  |  |
| --- | --- | --- | --- | --- | --- | --- | --- | --- | --- | --- | --- |
|  |  | Training dataset (cross-validation) |  |  |  |  | Validation/Independent dataset |  |  |  |  |
| Model | Thr | Sens | Spec | Acc | AUC | MCC | Sens | Spec | Acc | AUC | MCC |
| **DT** | 0.51 | 54.12 | 60.57 | 57.35 | 0.573 | 0.147 | 65.22 | 44.29 | 54.68 | 0.548 | 0.097 |
| **RF** | 0.52 | 67.38 | 69.89 | 68.64 | 0.745 | 0.373 | 68.12 | 67.14 | 67.63 | 0.735 | 0.353 |
| **LR** | 0.51 | 65.23 | 65.95 | 65.59 | 0.682 | 0.312 | 73.91 | 57.14 | 65.47 | 0.688 | 0.315 |
| **XGB** | 0.57 | 67.38 | 67.03 | 67.20 | 0.734 | 0.344 | 73.91 | 65.71 | 69.78 | 0.721 | 0.398 |
| **KN** | 0.50 | 66.31 | 62.72 | 64.52 | 0.712 | 0.291 | 85.51 | 50.00 | 67.63 | 0.723 | 0.379 |
| **GNB** | 0.51 | 63.08 | 62.72 | 62.90 | 0.681 | 0.258 | 69.57 | 58.57 | 64.03 | 0.672 | 0.283 |
| **SVC** | 0.51 | 66.67 | 64.87 | 65.77 | 0.735 | 0.315 | 78.26 | 64.29 | 71.22 | 0.760 | 0.429 |
| **ET** | 0.51 | 65.95 | 68.46 | 67.20 | 0.763 | 0.344 | 71.01 | 70.00 | 70.50 | 0.769 | 0.410 |

**Table 4:**
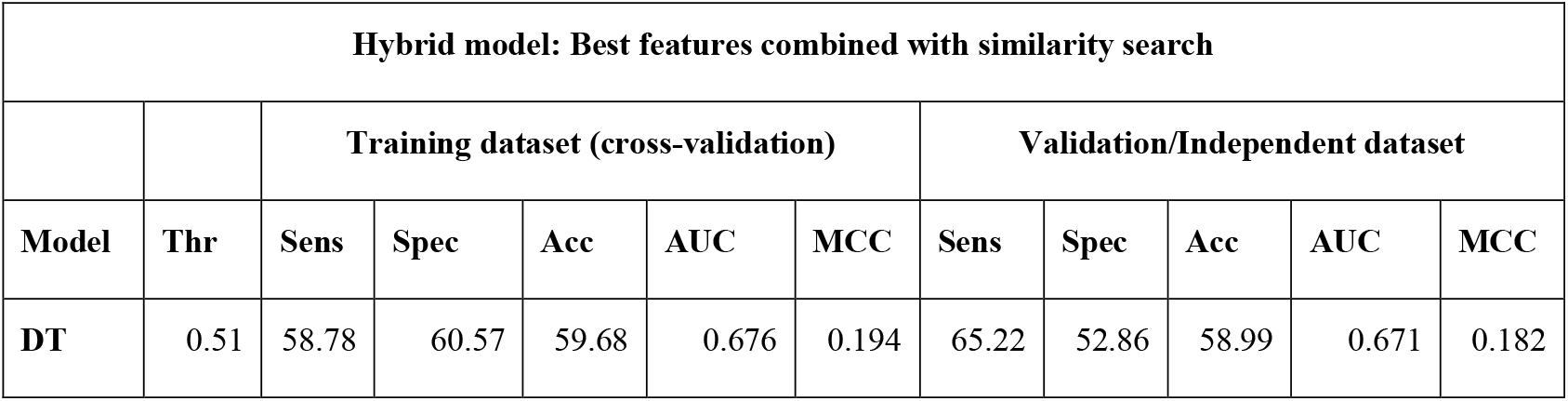
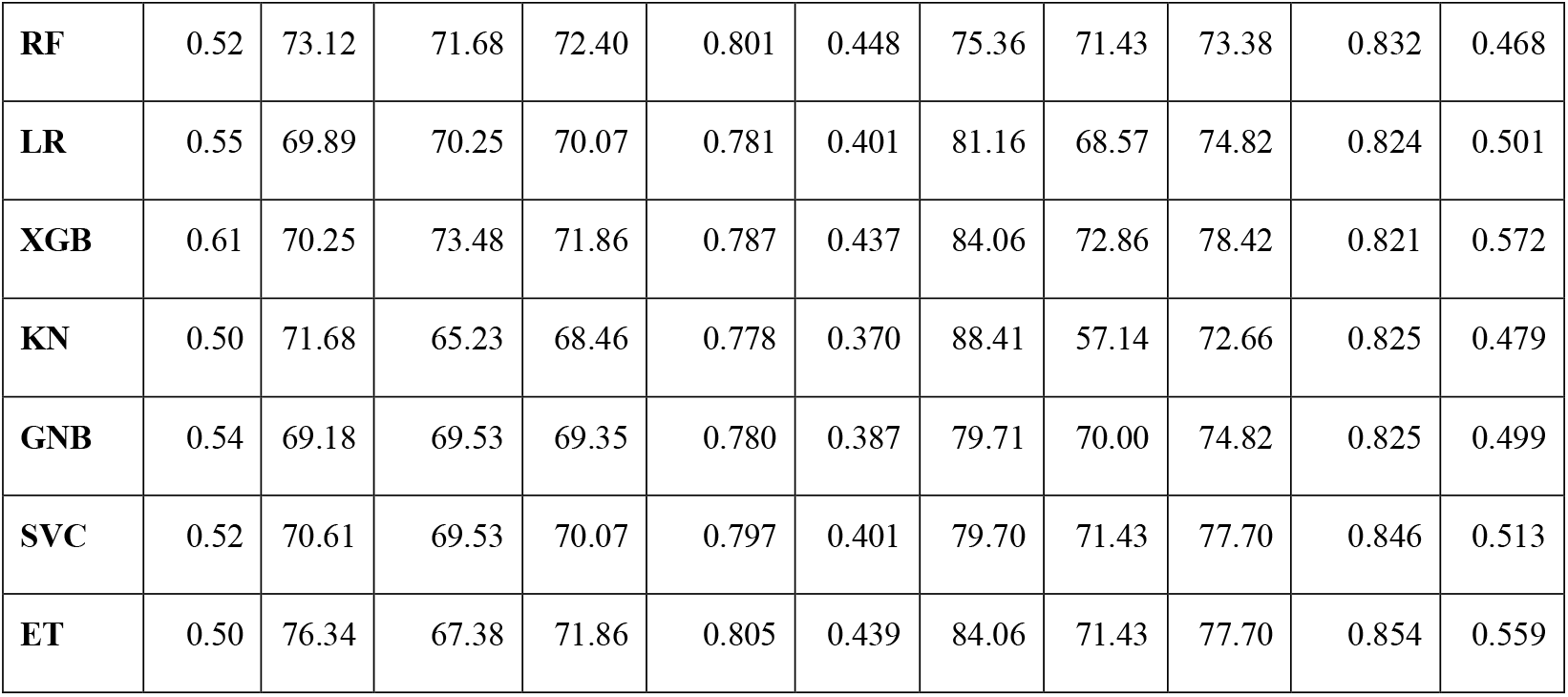
Results for hybrid model: best-performing features (AI-based methods) combined with similarity search (Alignment-based methods)

**Figure 2:**
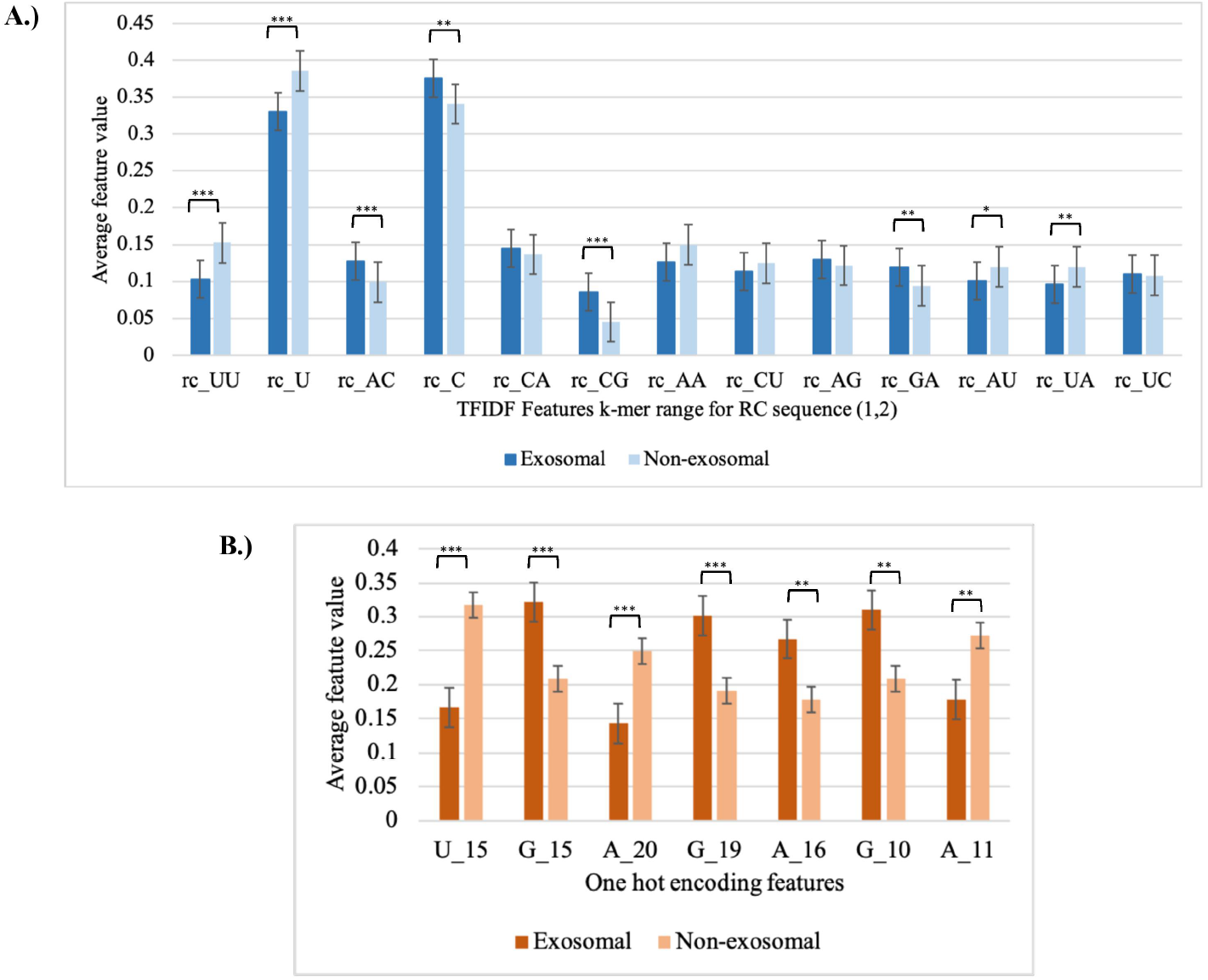
The top 20 most important features for the classification of exosomal (abundant in exosomes) and non-exosomal (non-abundant in exosomes) miRNA

**Figure 3:**
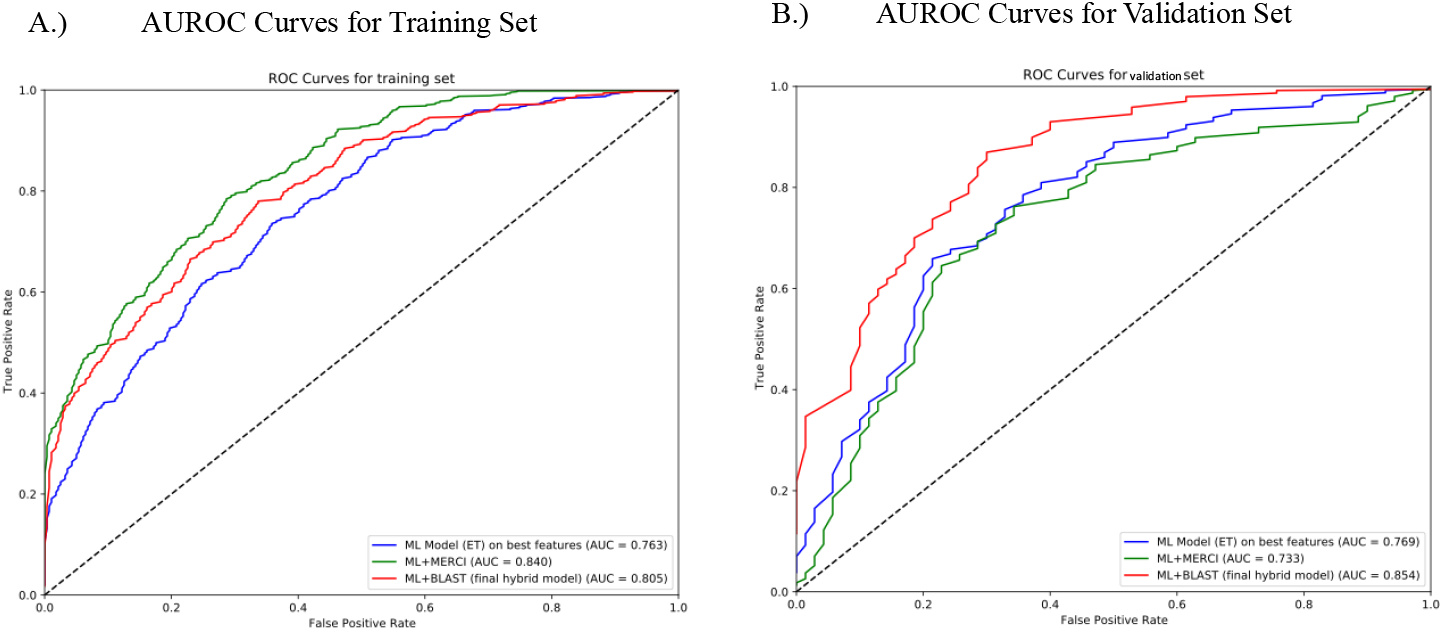
AUROC graphs for the best performing ML model, and hybrid models (ML + MERCI, ML + BLAST), for A) training set and B) validation set

### 3.4. Comparison with existing methods

We compared our prediction to the existing method – EmiRPred, which predicts exosomal miRNA and non-exosomal miRNA without taking expressions of different miRNA in exosomes into account [45]. In addition to this, we also compared our method against miRNALoc, and EL-RMLocnet that predict the subcellular localization of miRNA, with “exosome” being one of the possible locations [17,46]. To evaluate and compare the results, we input our independent validation set into the prediction server of miRNALoc that achieved an AUC of 0.422, compared to our server AdmirePred demonstrated a significantly higher AUC of 0.854. However, obtaining predictions from EL-RMLocNet was not feasible, as it processes only one sequence at a time, making it impractical to analyze the entire validation set of 139 sequences. As EmiRPred’s training dataset had a lot of the same miRNA used in AdmirePred, we found the miRNA sequences that were exclusively present in admirepred dataset and put them as input in EmiRPred server, it was able to predict the miRNA as exosomal and non-exosomal with an AUC of 0.623.

### 3.5. Webserver and Standalone Software

We wish to promote the field of non-invasive diagnostics and prognostics, focusing on miRNA based biomarkers and help the researchers worldwide working in this area. The developed web server offers three key modules: Predict, Design, and BLAST-search. The Predict module allows users to input query sequences and obtain predictions for abundant miRNA in exosomes. The Design module enables users to generate all possible mutant miRNA sequences and predict how likely a mutant miRNA is to be abundantly present in exosome. The BLAST-search module provides an option of similarity search against a database of known abundant and non-abundant miRNA sequences in exosomes. This web server is designed to be accessible across multiple devices, including smartphones, PCs, iMacs, and tablets. Additionally, we have developed a Python package, a standalone tool, and a GitHub repository, which are available at the following links: https://webs.iiitd.edu.in/raghava/admirepred/ and https://github.com/raghavagps/admirepred.

## 4. Discussions

Exosomal microRNAs (miRNAs) have emerged as critical players in intercellular communication and disease pathogenesis, offering promising potential as diagnostic and therapeutic biomarkers [5,47]. These small non-coding RNAs are selectively packaged into exosomes, which protect them from degradation and facilitate their transport to target cells, where they modulate gene expression [48]. Studies have demonstrated the utility of exosomal miRNAs in a variety of diseases, including cancer, neurodegenerative disorders, and cardiovascular diseases. For instance, in cancer, exosomal miR-21 and miR-155 have been identified as key regulators of tumor progression and metastasis, with elevated levels observed in the exosomes of patients with breast and pancreatic cancers [49,50]. In neurodegenerative diseases such as Alzheimer’s, exosomal miR-125b and miR-146a have been implicated in neuroinflammation and neuronal dysfunction [51]. Similarly, in cardiovascular diseases, exosomal miR-1 and miR-133a have been associated with myocardial injury and heart failure [52]. The ability of exosomal miRNAs to reflect disease states and their stability in bodily fluids make them attractive candidates for non-invasive diagnostics and targeted therapies.

To help accelerate exosome-based diagnostics and therapies, we conducted this research in predicting the miRNA present in abundance in exosomes during normal conditions. We believe this will help the scientists worldwide to compare the expression levels of miRNA inside exosomes during various disease conditions. To achieve this, we firstly employed alignment-based approaches including motif-search and similarity-search. We performed a motif-search to find motifs that are exclusively present in positive dataset (exosome abundant miRNA) for various conditions such as Gap = 0, 1, and 2. However, it was observed that this approach did not work well for the dataset as it had low coverage of about 7.6% and high error rate. For similarity-search approach, we used blastn-short. This method showed about 0% error but low coverage accounting to only about 34.7% of the total independent validation set sequences at e-value 10^−2^.

To overcome the limitations of alignment-based methods, we performed built machine learning-based models to predict highly abundant miRNA in exosomes. We calculated a number of features of the miRNA sequences including binary-profile based features and composition-based features like nucleotide composition, entropy, correlation, term-frequency-inverse document frequency (tfidf) etc. The best performing based features were binary-profile based features and tfidf features for reverse complementary sequences for k-mers (1,2). After combining these best-performing features, we achieved a maximum AUC of 0.769 and MCC of 0.410 on independent validation dataset. Next, we employed a hybrid method to harness the predicting power of both alignment-based and machine-learning based classification methods. After combining these techniques, we obtained the highest AUC of 0.854 and MCC of 0.559.

We compared our method against existing computational approaches designed to predict the subcellular localization of miRNAs, including those that specifically account for exosomal miRNAs, such as EmiRPred, EL-RMLocnet and miRNALoc [17,45,46]. These methods have been widely used to infer the localization of miRNAs within cellular compartment. However, our method demonstrated superior performance in accurately predicting miRNA localization, particularly in identifying abundant miRNAs in exosomes. This improvement can be attributed to our incorporation of miRNA expression profiles. We believe that the expression levels of miRNAs play a critical role in determining their abundance. For instance, miRNAs that are highly expressed in specific cell types or under certain disease states are more likely to be selectively packaged into exosomes, as exosomal sorting mechanisms often prioritize miRNAs that are functionally relevant to intercellular communication. By integrating expression data into our predictive framework, we were able to capture these nuances, leading to more accurate predictions.

## Author’s contributions

AA collected and processed the data, implemented the algorithms, developed the prediction models, and built the front end and back end of the web server. AA, and GPSR prepared the manuscript. GPSR conceived and coordinated the project. All authors have read and approved the final manuscript.

## Acknowledgements

The authors are thankful to the Council of Scientific and Industrial Research (CSIR) for providing the fellowship. The authors are also thankful to the Department of Computational Biology, IIITD, New Delhi for its infrastructure and facilities. We thank the Department of Biotechnology (DBT) for providing an infrastructure grant to the institute (Grant BT/PR40158/BTIS/137/24/2021). We would like to acknowledge that figures were created using BioRender, and English was corrected using Grammarly.

## Data Availability Statement

All the datasets generated in this study are available at https://webs.iiitd.edu.in/raghava/admirepred/dataset.php and codes are available at https://github.com/raghavagps/admirepred.

